# Local PI(4,5)P_2_ generation controls fusion pore expansion during exocytosis

**DOI:** 10.1101/2022.01.26.477824

**Authors:** Muhmmad Omar-Hmeadi, Alenka Guček, Sebastian Barg

## Abstract

Phosphatidylinositol(4,5)bisphosphate (PI(4,5)P_2_) is an important signaling phospholipid that is required for regulated exocytosis and some forms of endocytosis. The two processes share a topologically similar pore structure that connects the vesicle lumen with the outside. Widening of the fusion pore during exocytosis leads to cargo release, while its closure initiates kiss&run or cavicapture endocytosis. We show here, using live cell TIRF microscopy of insulin granule exocytosis, that transient accumulation of PI(4,5)P_2_ at the release site recruits components of the endocytic fission machinery, and stalls the late fusion pore expansion that is required for peptide release. The absence of clathrin differentiates this mechanism from clathrin-mediated endocytosis. The PI(4,5)P_2_ transients result from local phosphatidylinositol-phosphate-5-kinase-1c (PIP5K1c) activity, and knockdown of PIP5K1c, or optogenetic ablation of PI(4,5)P_2_ promotes fusion pore expansion. Thus, local phospholipid signaling controls fusion pore expansion peptide release through an unconventional endocytic mechanism.

## Introduction

Regulated exocytosis is a fundamental cellular mechanism that allows timely release of hormones, neurotransmitters and proteins. For example, pancreatic β-cells use exocytosis to release insulin that is stored in secretory granules during periods of elevated blood glucose. Exocytosis fuses the secretory vesicle membrane with the plasma membrane, starting with the formation of a narrow fusion pore (diameter 1-3 nm) (Lindau and Alvarez De Toledo, 2003) that establishes aqueous contact between vesicle lumen and extracellular medium. Initially, opening of the fusion pore allows small molecules and ions to be released, while the larger peptide hormones are retained in the vesicle (Obermuller et al., 2005). Similar to an ion channel, the fusion pore can flicker between open and closed states, before it eventually widens enough to allow peptide release. The empty vesicle then either flattens into the plasma membrane (*full fusion*) or is retrieved essentially intact by an endocytosis-like mechanism before complete mixing of the two membranes occurs (*kiss&run* or *cavicapture* (Obermuller et al., 2005; Taraska et al., 2003; Tsuboi et al., 2004). Fusion pore lifetimes vary greatly between systems, depending on vesicle size, cargo and other factors, and range from a few hundred milliseconds in most dense core granules to minutes for surfactant release (Dietl et al., 2001). Both kiss&run exocytosis and extended fusion pore lifetimes allow separate release of small neurotransmitters and larger peptide hormones from the same vesicle (Obermuller et al., 2005). While there is good evidence that fusion pore behavior is physiologically relevant (Barg and Gucek, 2016; Collins et al., 2016), the mechanisms and regulation of fusion pore behavior are poorly understood.

In principle, the behavior of the fusion pore may be controlled by promoting either its expansion, e.g. by SNARE proteins (Wu et al., 2017), synaptotagmin (Das et al., 2020), and the cytoskeleton (Sokac et al., 2003), or by restricting and stabilizing its hourglass-shaped neck. The latter mechanism is reminiscent of the scission event that occurs during endocytic vesicle budding. Dynamin, a GTPase responsible for fission during clathrin mediated endocytosis (CME), is indeed found at sites of exocytosis and its presence prolongs fusion pore lifetime (Guček et al., 2019; Somasundaram and Taraska, 2018; Trexler et al., 2016). A number of other CME proteins are also present at the exocytosis site (Trexler et al., 2016) with the notable exception of the cage forming protein clathrin (Tsuboi et al., 2004). Several of these proteins, including the BAR-domain protein amphiphysin, contain membrane curvature sensitive domains or amphipathic helices that bind to the neck of the endocytic fission site, and are candidates for controlling fusion pore dynamics during exocytosis.

Both exocytosis and CME are regulated by PI(4,5)P_2_ (Gong et al., 2005; Milosevic et al., 2005; Posor et al., 2013), a negatively charged phosphoinositide (PI) that contributes to membrane identity and acts as important cellular signaling molecule. PI(4,5)P_2_ recruits cytosolic proteins to specific locations on the inner face of the plasma membrane, by means of their pleckstrin homology- (PH), ENTH, PX or C2-domains (Gambhir et al., 2004; Lemmon, 2003; Martin, 2012). PI(4,5)P_2_ is present at the release site (Aoyagi et al., 2005; Kabachinski et al., 2014; Martin, 2015; Shin et al., 2018; Trexler et al., 2016) and promotes release probability (priming) and establishment of the fusion pore (Gong et al., 2005; Milosevic et al., 2005) by facilitating membrane binding of the C2-domain containing proteins Munc13 and synaptotagmin (Das et al., 2020; Pinheiro et al., 2016). Moreover, the SNARE protein syntaxin can aggregate in presence of PI(4,5)P_2_, which in turn affects vesicle docking at the plasma membrane (Aoyagi et al., 2005; Gandasi et al., 2014; Honigmann et al., 2013; Kabachinski et al., 2014). During CME, PI(4,5)P_2_ is generated at clathrin coated pits and orchestrates the sequential recruitment of components of the fission machinery (Lampe et al., 2016). Furthermore, PI(4,5)P_2_ directly affects membrane curvature and may support the highly curved hemifusion intermediates during endo- and exocytosis (Chernomordik et al., 1995). PI(4,5)P_2_ is synthesized by consecutive action of two distinct but related PI-kinases (van den Bout and Divecha, 2009), first phosphorylation of phosphatidylinositol to phosphatidylinositol 4-kinase (PI4K), and second phosphorylation of the resulting PI(4)P by phosphatidylinositol 4-phosphate 5-kinase (PIP5K).

Since PI(4,5)P_2_ is important for both exocytosis and CME, we hypothesized that it may control fusion pore behavior by initiating a kiss&run related endocytic mechanism. Using live-cell total internal reflection fluorescence microscopy (TIRF) in insulin-secreting β-cells, we find that PI(4,5)P_2_ is acutely generated at the time and location of fusion pore formation. The transient PI(4,5)P_2_ that is localized with the fusing granule recruits several proteins of the endocytic fission machinery, and strongly affects fusion pore behavior. We propose that PI(4,5)P_2_ couples exocytosis to a dynamin-dependent mechanism that controls the efficiency and timing of peptide release and prepares the vesicle membrane for rapid retrieval.

## Results

### Phosphoinositides accumulate transiently during delayed peptide release

Acute depletion of PI(4,5)P_2_ strongly inhibits regulated exocytosis (Milosevic et al., 2005; Omar-Hmeadi et al., 2018), suggesting an important role of this phospholipid during late stages of exocytosis. To understand this role, we studied PI(4,5)P_2_ dynamics during stimulated exocytosis. Insulin-secreting INS1 cells expressing a fluorescent PI(4,5)P_2_ sensor (EGFP-PH-PLCδ1) together with the granule marker NPY-tdmOrange2 were imaged by TIRF microscopy (Fig 1A). When exocytosis was stimulated by depolarizing the cells with medium containing elevated K^+^ (75 mM), individual granules underwent exocytosis and released their content, seen as the sudden disappearance of fluorescence puncta of the granule marker (see examples in Fig 1B and video). Fluorescence of the PI(4,5)P_2_-marker was relatively uniform throughout the cells’ footprint and unrelated to the position of granules, as reported previously (Omar-Hmeadi et al., 2018). However, at most sites of exocytosis we observed transient local accumulation of the PI(4,5)P_2_ sensor that was diffraction limited in size (<200nm) and temporarily and spatially aligned with the exocytosis event (Fig 1A,B). We and others have previously shown that exocytosis of individual granules occurs with two phenotypes (Barg et al., 2002), where the labeled peptide content is either released immediately (*Immediate Release* or IR, Fig 1B example 3), or after a delay during which the signal transiently increases (*Flash Release* or FR; Fig 1B example 1-2). This flash results from dequenching of the pH-sensitive granule marker after fusion pore opening, as long as the pore has not expanded sufficiently to allow peptide release. The duration of the flash can therefore be used to quantify fusion pore lifetime (Fig 1C; (Guček et al., 2019)). Notably, accumulation of the PI(4,5)P_2_ sensor signal was observed almost exclusively during events with the flash phenotype (Fig 1B). Indeed, when we separated the two event types and aligned to the moment of release, the average PI(4,5)P_2_ -sensor signal increased strongly during the FR events (164 events in 24 cells, Fig 1D, black), compared to no accumulation at the IR events (83 events in 24 cells, Fig 1D, grey P=5E^−8^, t-test).

**Fig 1:**
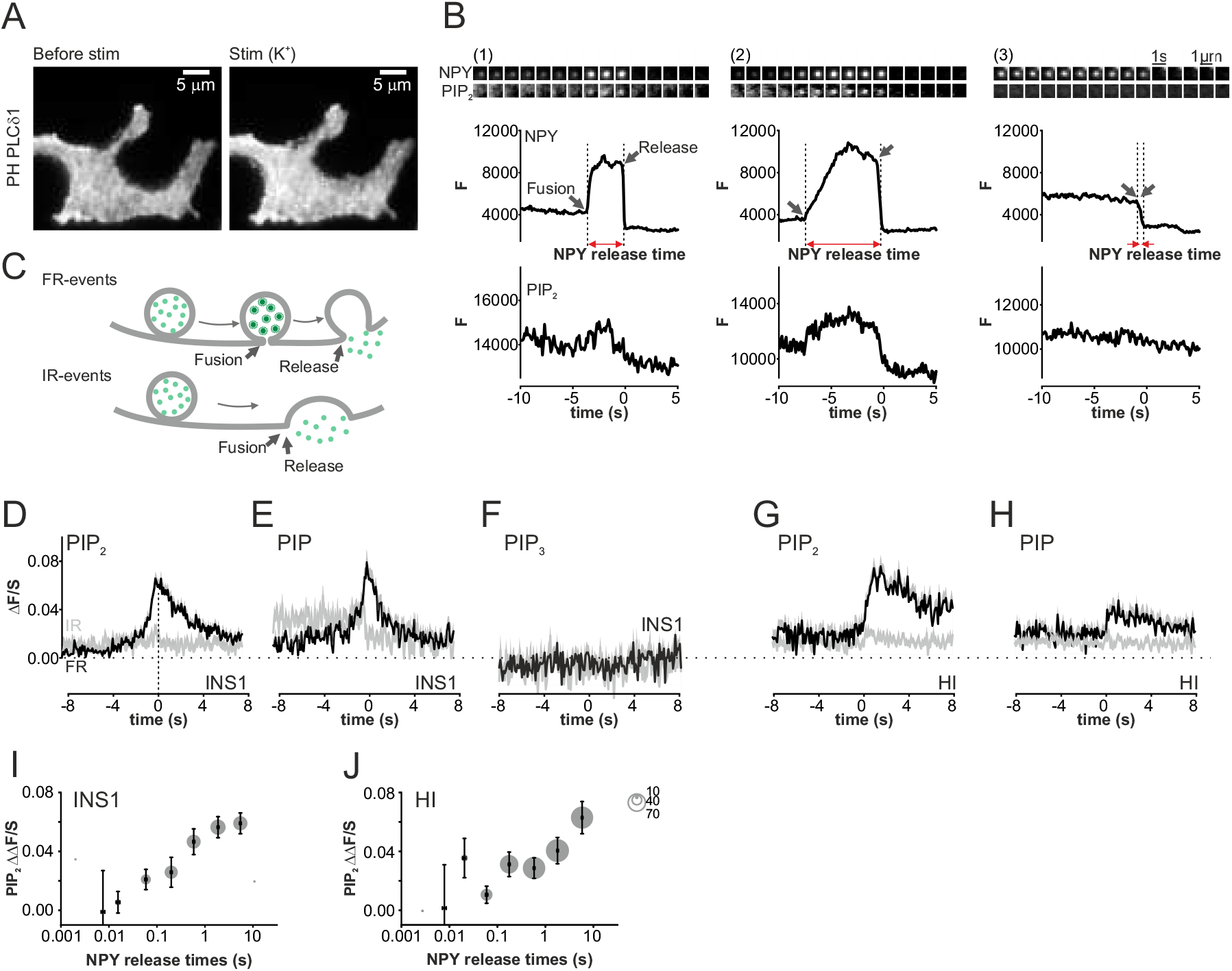
Transient accumulation of PI(4)P and PI(4,5)P_2_ at insulin granule exocytosis sites. (A) Representative example of an INS1 cell expressing the PI(4,5)P_2_ sensor PH-PLCδ1 in unstimulated conditions (max pixel intensities for 10 s before stimulation, left), and during stimulation with elevated K^+^ (max pixel intensities during first 20 s of stimulation, right). (B) Three examples (1-3) of single granule exocytosis in K^+^ stimulated cells, as in A. Image frames (upper) show granule (gr) and PI(4,5)P_2_ sensor (PIP_2_) images at the exocytosis site (1s/frame, scalebar 1μm), and traces the quantification of gr (middle) and PIP_2_ signal (lower) temporally aligned to the same events. Note temporary increase of gr fluorescence (indicating FR events with arrested fusion pore) and associated PI(4,5)P_2_-signal in examples 1-2, but not 3 (IR events). Black arrows indicate moment of fusion pore opening and content release, red the fusion pore lifetime. (C) Cartoons illustrating the interpretation of events in B. (D-F) Dynamics of phosphatidylinositol species at FR (black) and IR (grey) exocytosis events, quantified as average ΔF/S (see methods) for (D) the PI(4)P sensor EGFP-P4M-SidM (107 FR events and 84 IR events in 26 INS1-cells, 3 preps), (E) the PI(4,5)P_2_ sensor EGFP-PH-PLCδ1 (164 FR events, 83 IR events in 24 INS1-cells, 3 preps), and (F) the PI(3,4,5)P_3_ sensor 4EGFP-GRP1 (83 events in 15 INS1-cells, 3 preps). All signals were aligned to the moment of release (t=0); positive ΔF/S indicates excess sensor fluorescence at the exocytosis site. (G-H) As in D-F but for human islets cells, and aligned to the moment of fusion. (G) PI(4,5)P_2_ (228 FR events, 102 IR events in 33 cells), (H) PI(4)P (191 FR events, 234 IR events in 30 cells, 3 preps). (I) PI(4,5)P_2_ accumulation (ΔΔF/S) as function of fusion pore lifetime in INS1-cells (I) and human islet cells (J); same experiments as in D and H; circle size indicates events per bin.

Since local PI kinase activity may underlie the observed PI(4,5)P_2_ transients, we tested two other PIs. The PI(4)P marker (EGFP-P4M-SidM) also accumulated transiently at FR fusion sites (Fig 1E, black, 107 events, 26 cells), but not during IR events (grey, 84 events, P=4E^−8^, t-test). In contrast, the PI(3,4,5)P_3_ marker (4EGFP-GRP1) showed no peak-to-peak changes during either event type (83 events, 15 cells, Fig 1F, P=0.63, t-test). Moreover, acute PI accumulation during exocytosis was also observed in primary human β-cells (HI) expressing PH-PLCδ1-EGFP (228 FR events in 33 cells, 102 IR events, P=0.01, t-test, Fig 1G) or EGFP-P4M-SidM (30 cells, 191 FR events, 234 IR events, P=2.5E^−4^, t-test, Fig 1H), suggesting that the phenomenon is not species- or cell-type dependent.

We tested whether the amplitudes of individual PI(4,5)P_2_ transients correlate with fusion pore stability. The latter was estimated by measuring the time between fusion and release using a fitting procedure (NPY release time; see methods). The distribution of NPY release times followed a mono-exponential decay (Fig S1) and were on average 1.77 ± 0.15 s (247 granules in 24 cells). The release times correlated positively with the amplitude of the associated PI(4,5)P_2_ transient, for both INS1 cells (r_s_=0.33, Fig 1I) and human islet (r_s_=0.22 for HI, Spearman’s rank correlation, Fig 1J). The findings suggest that phosphoinositides are involved in the regulation of fusion pore dynamics.

### PI(4,5)P_2_ transients during FR exocytosis are not due to diffusion

The PI(4,5)P_2_ transients during exocytosis may result from either local synthesis or acute recruitment of the PI to the release site. We tested the latter by loading NPY-tdOrange2 expressing cells with a tail-labeled fluorescent PI(4,5)P_2_ analog (BODIPY FL-PI(4,5)P_2_) that enters the cytosolic leaflet of the plasma membrane (Omar-Hmeadi et al., 2018). The PI(4,5)P_2_ analog distributed near uniformly in the plasma membrane, and the granule associated signal (ΔF/S) of 0.001±0.0027 (87 events in 18 cells) indicated that there was no preference for granule sites. Importantly, ΔF/S did not change during exocytosis (Fig 2A, P=0.9, paired t-test), and labeled PI(4,5)P_2_ did therefore not accumulate at granules during these events. Redistribution or unmasking of PI(4,5)P_2_ during exocytosis as cause for the transients observed with PH-PLCδ1-EGFP (Fig 1B,D,G) can therefore be excluded.

**Fig 2:**
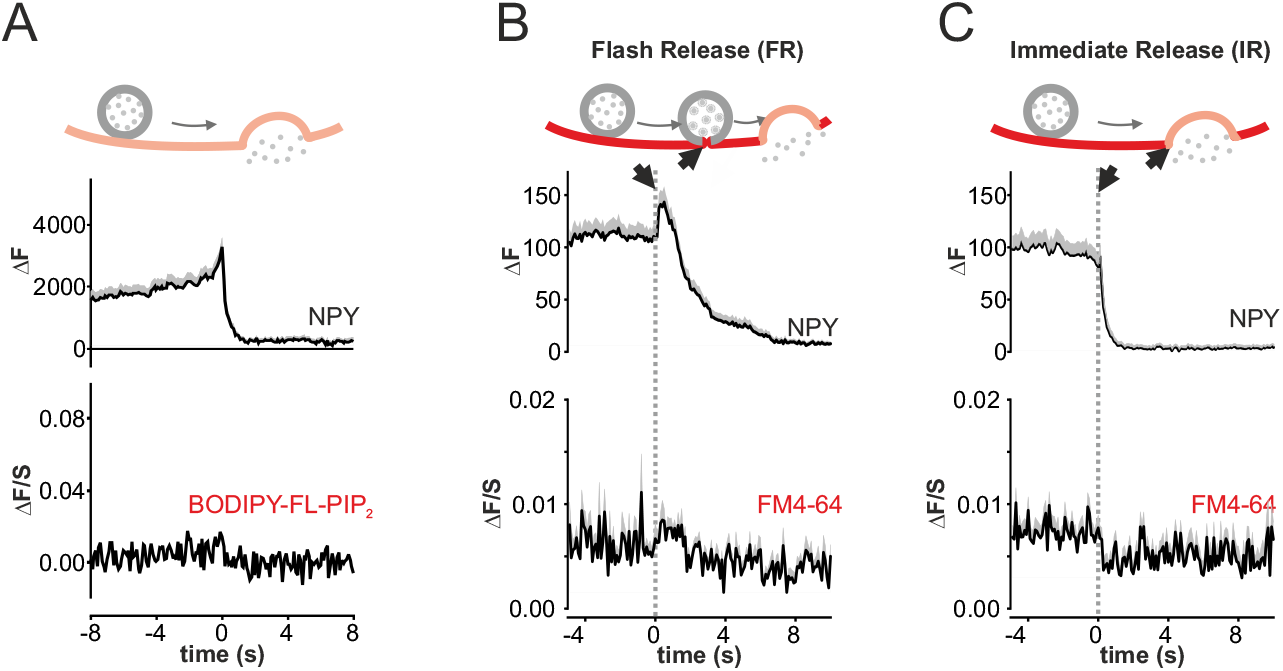
PI(4,5)P_2_ transients cannot be explained by lipid diffusion. (A) Average time course (+SEM) of NPY-tdmOrange2 (top) granule fluorescence in INS1 cells labelled with BODIPY FL-PI(4,5)P_2_ undergoing 75 mM K^+^-stimulated exocytosis (23 events, 8 cells, 3 preps). (B) Average time course (+SEM) of NPY EGFP (top) fluorescence events in INS1 cells labeled with FM 4-64 (5 μg/mL) 30 s before and during the K^+^-stimulated exocytosis for FR events (left, n=81) (C) and IR events (right, n= 56, 14 cells, 3 preps).

We also considered the possibility that the PI(4,5)P_2_ transients result from diffusion of PI(4,5)P_2_ into the granule’s membrane during exocytosis (Shin et al., 2018), which would appear as a local increase in PI(4,5)P_2_ before the granule has flattened into the plasma membrane. This effect was quantified by acutely saturating the plasma membrane, but not granules, with the lipophilic marker FM4-64 for 30 s, before stimulating exocytosis with elevated K^+^. In these conditions we did not observe any increase, but instead a small decrease in the granule-associated FM4-64 signal during exocytosis. The latter may result from the collapse of the unlabeled granule, causing displacement of labeled plasma membrane (Fig 2B-C). We conclude that PI(4,5)P_2_ transients at exocytosis sites do not result from local redistribution or unmasking.

### Delayed release during flash exocytosis is independent of clathrin

Phosphoinositides are well known to orchestrate the sequential recruitment of endocytic proteins during clathrin mediated endocytosis (CME) (Posor et al., 2013; Taylor et al., 2011), and several proteins involved in CME are also found at insulin granule sites (Trexler et al., 2016). We therefore quantified recruitment of two of these proteins, clathrin and amphiphysin, during exocytosis events (Fig 3). Clathrin light chain (CLC, labeled with GFP) was coexpressed with NPY-tdmOrange2 and exocytosis was stimulated by depolarization with elevated K^+^. CLC expression resulted in a punctate staining as expected for CME structures. Before exocytosis, the CLC signal did not colocalize with granules and its ΔF/S value was essentially zero. During FR exocytosis events (Fig 3A), the CLC signal remained zero at the release site (P=0.2, paired t-test), while IR events (Fig 3B) were associated with a modest and slow increase consistent with compensatory CME. In contrast, amphiphysin (as Amphiphysin1-mCherry, coexpressed with NPY-EGFP) was present at the release site before both FR and IR-exocytosis, and additional transient recruitment occurred during FR exocytosis (Fig 3A), but not during IR-exocytosis (Fig 3B, brown). The peak of this amphiphysin recruitment coincided with the moment of peptide release and with the peak of the PI(4,5)P_2_ transient detected using PH-PLCδ1-EGFP (Fig 3A). The data indicate that the mechanism that delays peptide release during FR exocytosis uses amphiphysin, but not clathrin, distinguishing it from classical CME.

**Fig 3:**
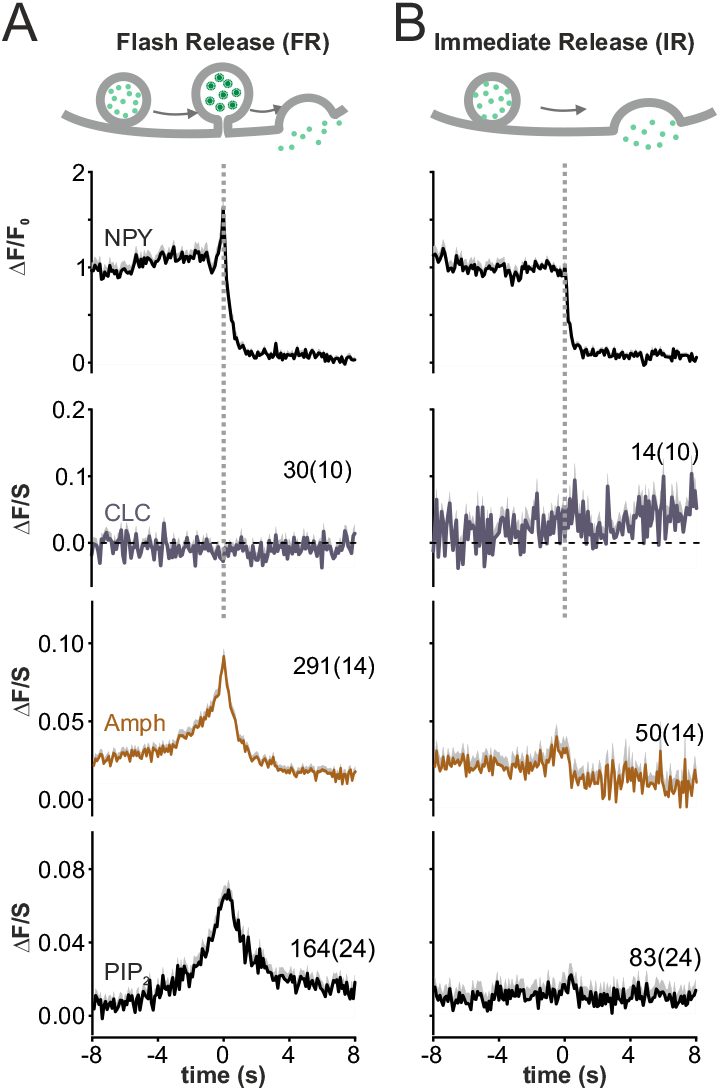
PI(4,5)P_2_ transients are distinct from clathrin mediated endocytosis. (A-B) Average time course (+SEM) of NPY-tdmOrange2 (black), GFP-CLC (blue), amphiphysin mCherry (brown) and the PI(4,5)P_2_ sensor EGFP-PH-PLCδ1 (black, from Fig 1) fluorescence during FR exocytotic events (A) or IR events (B). Dashed line indicates signal alignment from the same experiment. Numbers indicate event count (cell count).

### PI(4,5)P_2_ transients correlate with recruitment endocytosis-related proteins

Several endocytosis-related proteins are found at the insulin granule release site, and recruitment of dynamin to the exocytosis site prolongs fusion pore lifetime (Guček et al., 2019; Somasundaram and Taraska, 2018; Trexler et al., 2016). We hypothesized that PI(4,5)P_2_ orchestrates the recruitment of these proteins, and tested a panel of proteins with membrane binding domains for their ability to be recruited to the release site (Fig 4C). Limited by the two color channels of our microscope, we used PI(4,5)P_2_ transients (measured using PH-PLCδ1-EGFP) during K^+^-stimulation as proxy for FR exocytosis events (Fig 4A). This is reasonable, because we did not observe any PI(4,5)P_2_ transients in the 10 s before K^+^-stimulation (Fig S1E), indicating that a large majority of the observed events relate to exocytosis. During these events the PI(4,5)P_2_ signal reached its peak within ^~^1 s, before decaying back to baseline over the course of >10 s. Endocytosis-related proteins were individually co-expressed with the PI(4,5)P_2_ sensor (Fig 4B). Note that the PH-PLCδ1-EGFP does not saturate available PI(4,5)P_2_, because its expression has no effect on fusion pore behavior (Figs S1), and its binding to the plasma membrane increases with expression level (Omar-Hmeadi et al., 2018).

**Fig 4:**
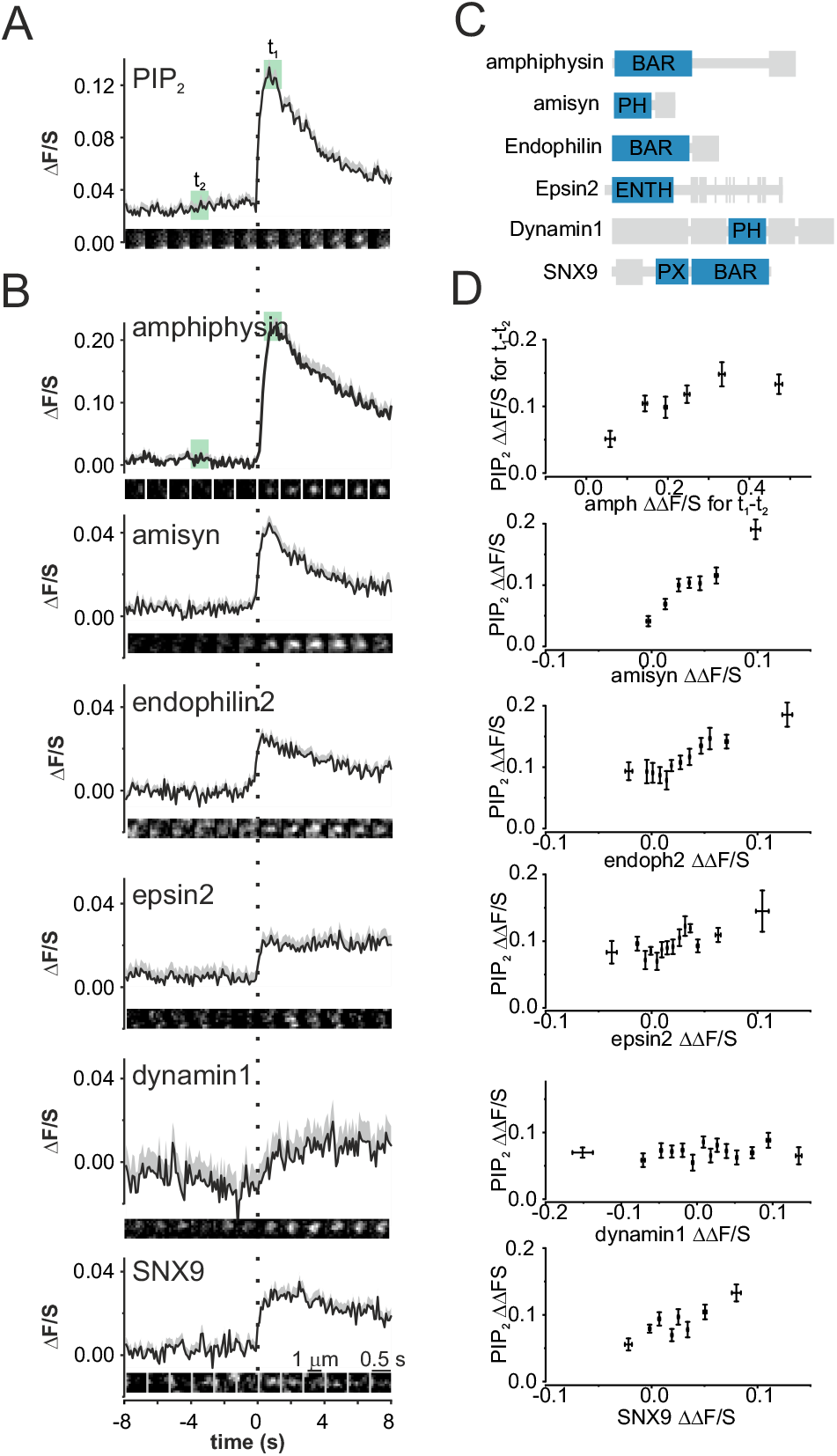
Endocytosis related proteins are recruited during K^+^-stimulated PI(4,5)P_2_ transients. (A) Average time course of PH-PLCδ1-EGFP transients (as proxy for FR exocytosis), during K^+^ stimulated exocytosis; same cells as in B (ΔF/S±SEM; n=79 events, 9 cells/2 preps). Image sequence shows an example event in amphiphysing co-expressing cells. (B) Associated average protein signal spatio-temporally aligned to events as in A, for cells co-expressing PH-PLCδ1-EGFP with either amphiphysin-mCherry (79 events, 9 cells/2 preps), mCherry-amisyn (102,13/3), endophilin2-mCherry (168 events, 18 cells /5 preps), epsin2-mCherry (199 events, 23 cells /3 preps), dynamin1-GFP (with PLCδ1-mRFP, 216 events,33 cells /7 preps), or mCherry-SNX9 (122 events,15 cells /3 preps). Image sequences shows example events. (C) Schematic illustration of protein domains for proteins used in A-B, with PI(4,5)P_2_ binding regions in blue. (D) Peak amplitudes of PH-PLCδ1-EGFP (ΔΔF/S for t_1_-t_2_ in A) plotted as function of peak amplitudes of proteins in B. Each symbol represents average (±SEM).

Both visual inspection and quantitative analysis (Fig 4B,D) revealed that the PH-domain containing proteins amisyn (P=2.3E-15), EGFP-dynamin1 (P=0.71), the ENTH domain protein epsin2 (P=0.008), the PX domain protein SNX-9 (P=2E-6) and the BAR domain proteins amphiphysin (P=3.9E-32) and endophilin2 (P=0.16, paired t-tests; Fig 4B) were all recruited to the site of exocytosis-associated PI(4,5)P_2_ transients. Most of the proteins mirrored the <1 s timecourse of the PI(4,5)P_2_ transient, with the exception of dynamin which was recruited slower. Amphiphysin, amisyn, endophilin and SNX9 signals decayed rapidly and in synchrony with the PI(4,5)P_2_ signal, while epsin2 and dynamin remained elevated at the release site for >8 s. For most of the proteins the recruitment to the release site correlated well with the amplitude of the PI(4,5)P_2_-sensor signal (both measured as ΔΔF/S between t_1_ and t_2_ in Fig 4A, which estimates the change in apparent affinity to the release site, as in Fig 4A). Dynamin1 was exceptional in that its recruitment was not affected by the amplitude PI(4,5)P_2_ transient. The data suggest that local PI(4,5)P_2_ signals directly drive the recruitment of a cluster of proteins to the exocytosis site, which in turn may lead to fusion pore restriction.

### Delayed release exocytosis depends on PI(4,5)P_2_

To investigate the role of PI(4,5)P_2_ in fusion pore regulation, we acutely depleted PI(4,5)P_2_ by recruiting phosphatidylinositol 5-phosphatase to the plasma membrane using an optogenic approach that exploits light-induced CRY2–CIBN dimerization. Phosphatidylinositol 5-phosphatase was fused to the soluble blue-light receptor CRY2 (CRY2-5-Ptase), and CRY2’s binding partner CIBN was anchored in the plasma membrane using the CAAX motif (CIBN-CAAX, Fig 5A). As expected, exposure to blue light caused rapid recruitment of GFP-Cry2-5Ptase to the plasma membrane followed by depletion of plasma membrane PI(4,5)P_2_ in cells co-expressing the two proteins with the PI(4,5)P_2_ sensor (Fig 5B).

**Fig 5:**
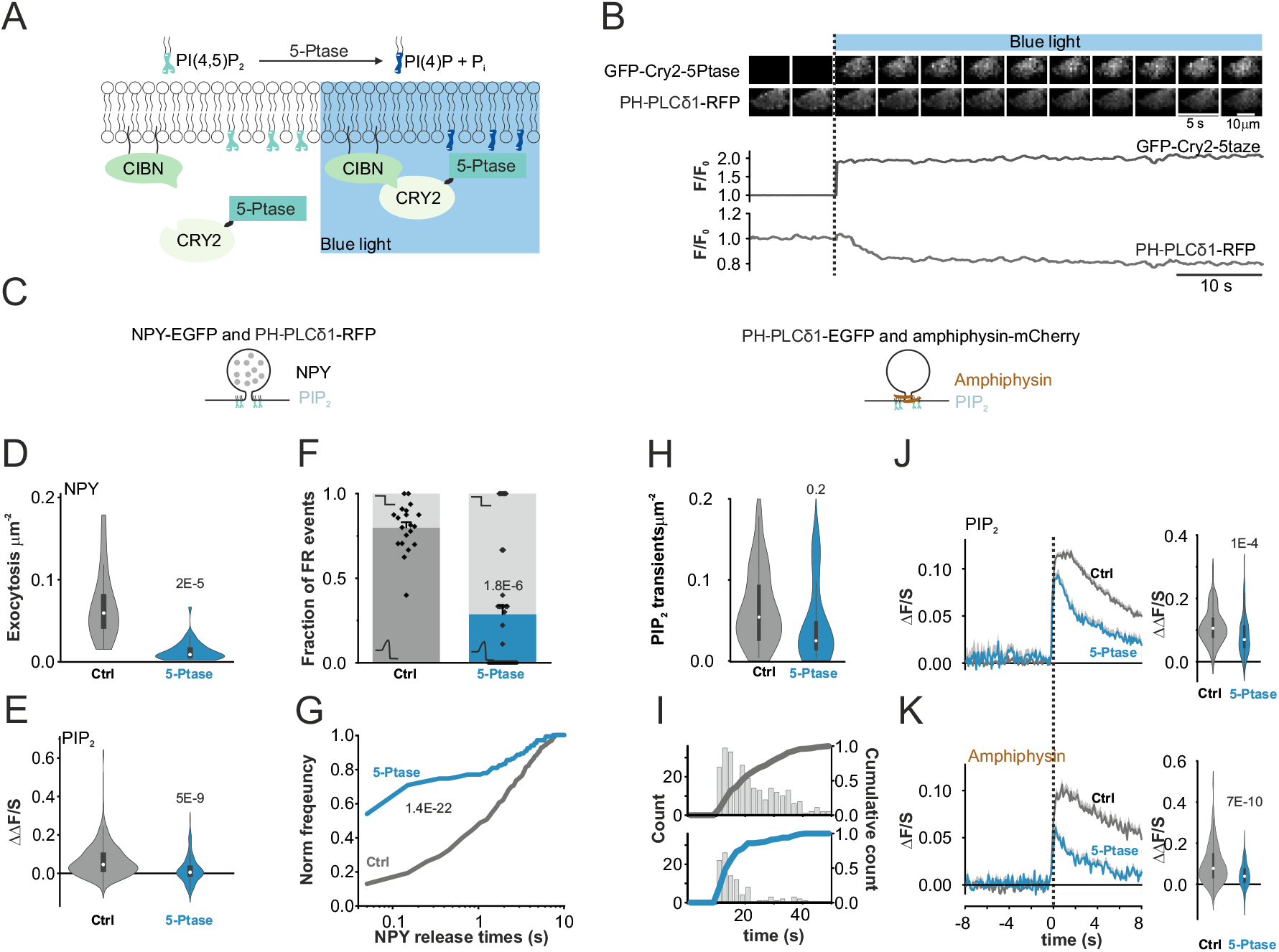
Acute optogenetic PI(4,5)P_2_ depletion impairs exocytosis but accelerates peptide release. (A) Cartoon illustrating the optogenetic Cry2-5-Phosphatase recruitment to the CIBN membrane anchor upon blue light activation and subsequent acute PI(4,5)P_2_ depletion. (B) Representative image example (top) and fluorescence time course (bottom) for an INS1 cell expressing GFP-Cry2-5Ptase and PH-PLCδ1-RFP. Note acute GFP-Cry2-5Ptase fluorescence increase upon blue light activation and subsequent decrease in PH-PLCδ1-RFP fluorescence. (C) Cartoons illustrating labeling strategies for experiment in D-G (left) and experiment in H-K (right). (D) Exocytosis during 40 s of K^+^-stimulation in cells co-expressing NPY-EGFP and PH-PLCδ1-RFP with either CAAX alone (Control, 20 cells, 3 preps) or both CAAX and Cry2-5Ptase (5-Ptase, 46 cells, 3 preps). (E) Distribution of PH-PLCδ1-RFP peak amplitudes (ΔΔF/S) in D. n=330 events (control), 130 events (5-Ptase). (F) Fraction of FR events in (D). (G) Cumulative histograms of NPY release times for events in D. P=1.4E-22, KS test (H) Frequency of PH-PLCδ1-EGFP transients during during 40 s of K^+^-stimulation in cells co-expressing PH-PLCδ1-EGFP and amphiphysin-mCherry with either CAAX alone (Control, 18 cells, 3 preps) or both CAAX and Cry2-5Ptase (5-Ptase, 14 cells, 3 preps). (I) Time course of PI(4,5)P_2_ transients for cells in H, for control (upper, black) and with Cry2-5Ptase (lower, blue). (J-K) Average time course (ΔF/S ±SEM) and distribution of peak amplitudes (ΔΔF/S, right panel) of EGFP-PH-PLCδ1 (J) or amphiphysin-mCherry in control (grey, 261 events, 18 cells, 3 preps), or with Cry2-5Ptase ( blue, 32 events, 38 cells, 3 preps).

Using this system, we assessed the effect of acute PI(4,5)P_2_ depletion on exocytosis in cells that additionally expressed NPY-EGFP as granule marker (Fig 5C). After 10 s exposure to blue light, K^+^-stimulated exocytosis was strongly reduced in cells expressing the complete optogenetic system (0.013 ± 0.001 μm^−2^ events per 40 s, 46 cells, 3 preps, blue, Fig 5D), compared to a control group in which the CRY2-5-Ptase was omitted (0.068 ± 0.01 μm^−2^ events per 40 s, 20 cells, 3 preps, grey, Fig 5D). PI(4,5)P_2_ transients had on average smaller amplitudes (Fig 5E, P=5E-9, t-test), and both the ratio of FR to IR exocytosis (Fig 5F, P=1.8E-6, u-test) and NPY release times (Fig 5G, P=1.4E-22, KS-test) were strongly decreased in cells expressing CRY2-5-Ptase, compared with control cells that did not. None of the parameters was affected by expressing CRY2-5-Ptase or CIBN-CAAX alone (Fig S5).

Next, we quantified the recruitment of amphiphysin during K^+^-stimulated exocytosis (Fig 5H-K). In cells expressing amphiphysin-mCherry, PI(4,5)P_2_ sensor, and the complete optogenetic system, blue light exposure decreased the frequency of exocytosis-related PI(4,5)P_2_ transients, compared with control cells in which CRY2-5-Ptase was omitted (Fig 5H-I); this effect was strongest during the latter half of the stimulation (Fig 5I, P=0.02, t-test). The amplitudes of both PI(4,5)P_2_- and amphiphysin-mCherry transients were reduced by about half when CRY2-5-Ptase was included, compared with control lacking the enzyme (Fig 5J-K). In both groups, the amphiphysin signal at the release site had a timecourse very similar to that of PI(4,5)P_2_. The data indicate that flash/DR exocytosis depends on localized PI(4,5)P_2_ transients, and confirm the general requirement of PI(4,5)P_2_ for granule priming.

### PIP5K activity is required for delayed release exocytosis

We tested the hypothesis that PI(4,5)P_2_ is generated locally by PIP5-kinase (PI5K) activity that phosphorylates PI(4)P to PI(4,5)P_2_ (Fig 6A) (Doughman et al., 2003). The most abundant isoform in INS1 cells, PIP5K1c (Blanchet et al., 2015), was targeted for knockdown (KD) using siRNA. Quantitative RT-PCR confirmed a ^~^90% reduction of PIP5k1c mRNA in cells treated with siRNA specifically targeting this isoform, compared with a scrambled siRNA as control (Fig 6B). Exocytosis and PI(4,5)P_2_ dynamics were then analyzed in PIP5K1c-KD and control INS1-cells. K^+^-stimulated exocytosis was strongly reduced in the KD cells compared with scrambled control (Fig 6C), as is expected for cells with overall reduced PI(4,5)P_2_-levels (Milosevic et al., 2005; Omar-Hmeadi et al., 2018; Xie et al., 2016). In the KD, the fraction of FR exocytosis events was reduced by on-third (Fig 6D), and NPY release times were decreased by 40% (Fig 6E; 2.5±0.2 s in ctrl vs 1.6±0.2 s in KD, P=4.6E^−4^, KS-test). PI(4,5)P_2_ transients were still observed in the KD cells, but their amplitude was reduced to one-third compared with scrambled siRNA control cells (Fig 6G; P=0.007, t-test). Since amphiphysin is recruited to PI(4,5)P_2_-enriched membranes (Somasundaram and Taraska, 2018), we also investigated how PIP5k1c knockdown affects amphiphysin behavior. Expression of amphiphysin-mCherry had no effect on exocytosis frequency, but slightly increased the fraction of FR events in scramble control cells; there was no additional effect of amphiphysin in the knockout (Fig 6C-D). Amphiphysin-mCherry accumulated at exocytosis sites in both groups, regardless of whether exocytosis was detected using NPY-EGFP (Fig 6H) or EGFP-PH-PLCδ1 (Fig S6). The results indicate that acute PI(4,5)P_2_ synthesis by PIP5K at the granule site is required for delayed release exocytosis.

**Fig 6:**
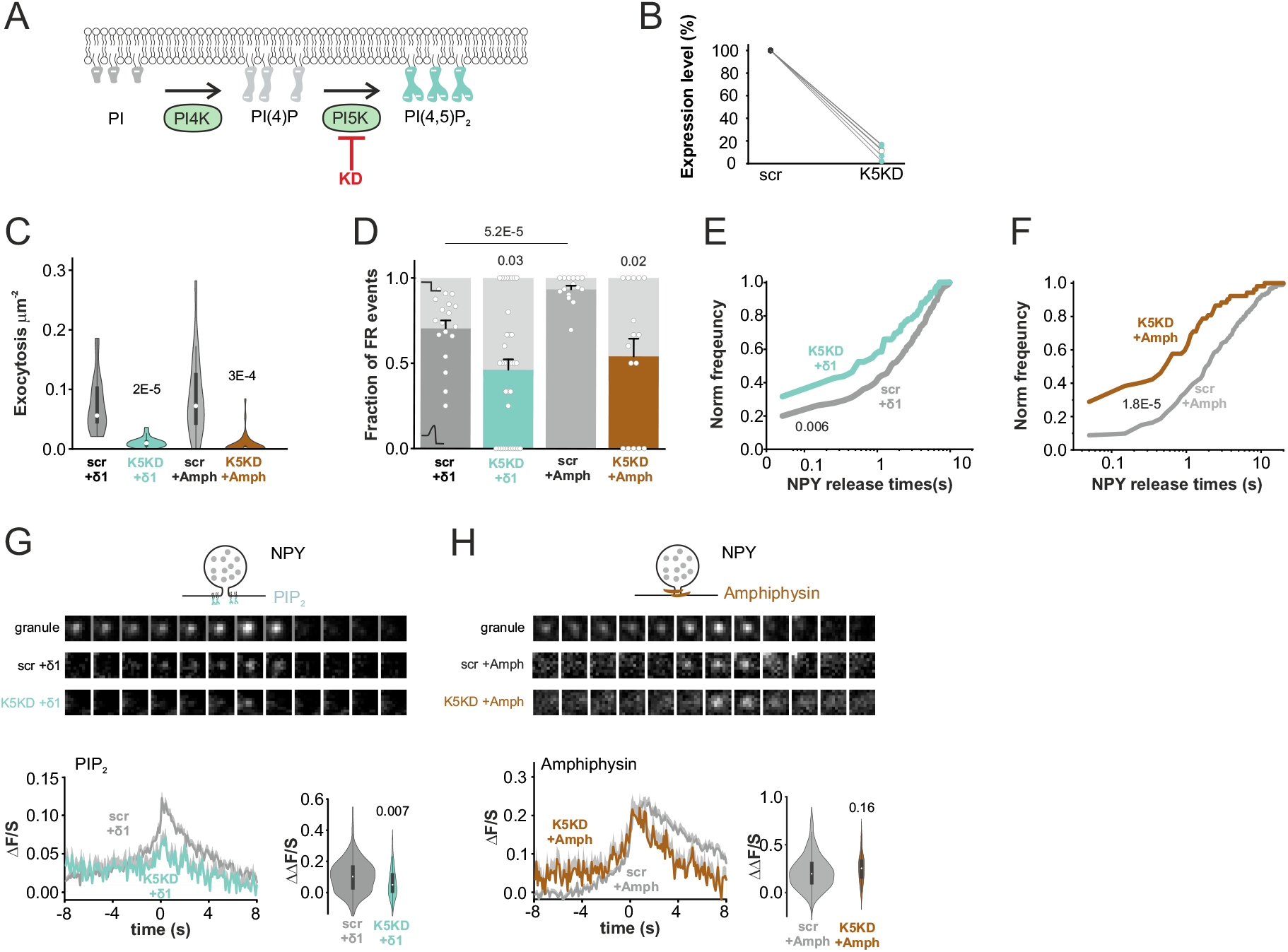
Knockdown of PI4P5-kinase accelerates peptide release. (A) Cartoon illustrating phosphatidylinositol synthesis steps. (B) Quantification of PIP5k1c expression in cells expressing scrambled control (scr) or PIP5k1c specific shRNA (K5KD). (C) Exocytosis during 40 s of K^+^-stimulation quantified in cells expressing NPY. The cells coexpressed combinations of scr, K5KD, EGFP-PH-PLCδ1, and amphiphysin-mCherry, as indicated (scr+PLCδ1, 18 cells; K5KD+PLCδ1, 38 cells; scr+amphiphysin, 16 cells; K5KD+amphiphysin, 38 cells. Values are p-values for scr vs K5KD, non-paired t-test. (D) Fraction of FR events in experiments in C, Values indicate p-values, u-test. (E) Cumulative frequency histograms of NPY release times in C, for scr+PH-PLCδ1 (grey) and K5KD+PH-PLCδ1 (light blue). P=0.006, KS test (F) As in (E) but for scr or K5KD + amphiphysin. P=1.8E-5, KS test. (G) Examples of single granule exocytosis (top), average time course of PH-PLCδ1 signal (ΔF/S ±SEM; lower left), and distribution of the peak amplitudes (ΔΔF/S, non-paired t-test; lower right) for scr+PH-PLCδ1 (grey; 210 FR events in 18 cells, 3 preps) and K5KD+PH-PLCδ1 (light blue; 40 FR events in 38 cells). Same experiments as in C-E. (H) As in (G) but for scr+amphiphysin (grey; 249 FR events, 16 cells) and K5KD+amphiphysin (brown; 32 events, 38 cells).

### PI4K is required for PI(4)P but not PI(4,5)P_2_ transients

Phosphatidylinositol 4-kinase (PI4K) maintains plasma membrane PI(4)P and thus indirectly levels of PI(4,5)P_2_ (Fig 7A). We therefore targeted PI4K2a activity for knockdown using a microRNA based approach, which leads to 60% reduction of its mRNA (Fig 7B). As expected, PI4K-knockdown or A1 treatment resulted in the elimination of PI(4)P transients that are seen in control during K^+^-stimulated exocytosis (P=8.5*10^−6^, oneway ANOVA, Fig 7H-L). Similar results were obtained when PI4K activity was instead blocked pharmacologically using the potent PI4K inhibitor A1 (100nM, P=6E-6, Fig 7J) (Bojjireddy et al., 2014); this effect was confirmed in primary human β-cells (Fig S7). Inhibition of PI4K activity by KD or A1 was accompanied by slightly smaller PI(4,5)P_2_ transients (Fig 7C-G) and a tendency to reduce exocytosis (Fig 7G,L), which is consistent with the fact that PI4K generates the substrate for PI(4,5)P_2_ synthesis. Treatment with A1 had no effect on the recruitment of amphiphysin-mCherry to the release site (Fig 7M,N,P), the fraction of FR events (Fig 7O), or NPY release times (Fig 7R). However, amphiphysin expression on its own increased both the ratio of FR events (Fig 7O) and NPY release times (Fig 7R, P=1.5E-11, KS-test), as observed earlier (Guček et al., 2019). We conclude that PI(4)P transients are not involved in the regulation of peptide release or the recruitment of amphiphysin to the release site.

**Fig 7:**
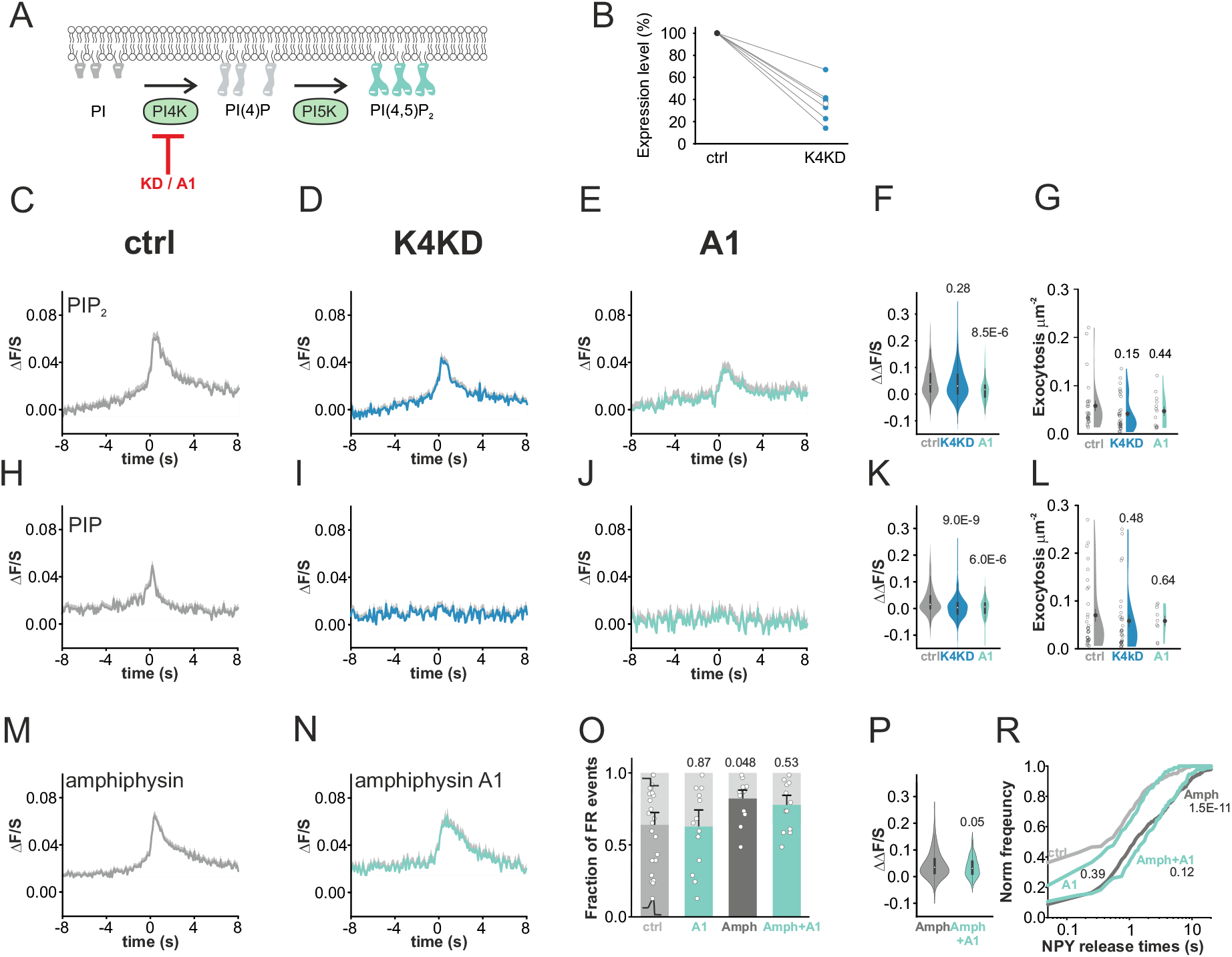
PI4-kinase inhibition prevents PI(4)P, but not PI(4,5)P_2_, transients. (A) Cartoon illustrating phosphatidylinositol synthesis. (B) Quantification of mirRNA based knockout of PI4K2a (K4KD) compared with control INS1 cells. (C-E) Average EGFP-PH-PLCδ1 time course (ΔF/S ±SEM) during K^+^-stimulation in cells co-expressing NPY-mOrange2; for ctrl (E, 225 events), K4KD (D, 291 events), or ctrl with kinase-4 inhibitor A1 (E, 110 events, 100nM A1 for 10 minutes). All events are spatiotemporally aligned to exocytosis events in the NPY-mOrange2 signal. (F) Distribution of peak amplitudes (ΔΔF/S ±SEM) for transients in C-E, p-values shown are for one-way ANOVA, Fisher posthoc test. (G) Exocytosis during 40 s K^+^-stimulation in C-E, for control (27 cells), K4KD (33 cells), and A1 (13 cells). p-values are for one-way ANOVA, Fisher posthoc test. (H-L) As in C-G, but for cells expressing EGFP-P4M-SidM in control (H, 264 events, 32 cells), K4KD (I, 245 events, 31 cells), or A1 (J, 103 events, 9 cells). (M-N) Average amphiphysin-mCherry time course for FR exocytosis events (ΔF/S ±SEM, aligned to rising phase of co-expressed NPYmOrange2 signal) in ctrl (M, 291 FR events) and with A1 (N, 142 FR events). (O) Fraction of FR events in cells expressing NPY-EGFP alone (C), or together with amphiphysin-mCherry (Amph), and in presence or absence of A1; ctrl (23 cells), control with A1 (13 cells), Amph (14 cells) or Amph with A1 (13 cells); p-values shown are for u-test. (P) Distribution of peak amplitudes (ΔΔF/S) for events in M-N (p-values for non-paired t-test). (R) Cumulative frequency histograms of NPY release times in (O). P=0.39 for ctrl vs A1, P=0.12 for Amph+A1 vs Amph, and P=1.5E-11 for Amph vs ctrl, KS-test).

## Discussion

We report here that exocytosis of insulin granules frequently coincides with locally generated PI(4,5)P_2_ transients at the release site, and that these events are associated with recruitment of the endocytic fission machinery and delayed peptide release. Since the latter reflects failure of the fusion pore to expand beyond a diameter that allows peptides to escape (Barg et al., 2002; Obermuller et al., 2005; Taraska et al., 2003; Tsuboi et al., 2004), we propose that PI(4,5)P_2_ acts as signal that coordinates a fission releated mechanism during exocytosis of dense-core granules. This mechanism is based on 1) spatial and temporal correlations of PI(4,5)P_2_ transients, pore lifetimes, and protein recruitment to the exocytosis site, 2) altered pore behavior in response to manipulations of protein and lipid levels, 3) the temporal sequence of PI(4,5)P_2_ and protein signals, and 4) the PI(4,5)P_2_ concentration dependence of the recruitment of most of the tested endocytosis proteins. Although we acknowledge that fusion pore size was not directly measured, our imaging methodology robustly detects the two physiologically relevant parameters of fusion pore behavior (initial pore opening, and its sudden expansion that permits hormone release) (Guček et al., 2019; Obermuller et al., 2005).

Since delayed fusion pore expansion temporally separates release of peptide hormones from that of smaller transmitter molecules (such as ATP or GABA) (Obermuller et al., 2005), it is not surprising that this mechanism is both regulated and physiologically relevant. cAMP-dependent signaling increases kiss&run exocytosis (MacDonald et al., 2006) through the cAMP-sensor Epac2 (REPGEF2) and delays insulin release in pancreatic β-cells (Guček et al., 2019). Interestingly, cAMP and PI(4,5)P_2_ share two opposing effects on peptide release; both delay fusion pore expansion, but also strongly activate priming and increase the number of readily releasable granules. Moreover, in type-2 diabetes over-expression of the SNARE binding protein amisyn (STXBP6) decreases insulin secretion by promoting kiss&run exocytosis (Collins et al., 2016) (but not in chromaffin cells (Kondratiuk et al., 2020)) and excess cholesterol causes insulin granules to fuse preferably through kiss-and-run mechanism (Xu et al., 2017). It can be speculated that by the mode of exocytosis secretory granules may serve different intercellular communication needs, separating hormonal and paracrine secretion depending on situation and perhaps intracellular location.

Presence of PI(,5)P_2_ in the plasma membrane promotes granule priming, and has been shown to shorten the foot signal in amperometric recordings (Gong et al., 2005; Milosevic et al., 2005). The latter reflects initial fusion pore behavior and the finding suggests that PI(,5)P_2_ promotes the transition from a brief pore flickering phase to a stable open state that allows release of small molecules, but not yet peptides (Guček et al., 2019). During this period of a few milliseconds, pore behavior is likely dictated by actions of the exocytosis machinery, since synaptobrevin (Ngatchou et al., 2010) and synaptotagmin/PI(4,5)P_2_ interaction promote the transition (Das et al., 2020). In contrast, the peptide release times observed optically here and elsewhere (Barg et al., 2002; Taraska et al., 2003) exceed by far the duration of the initial pore flickering (Lollike et al., 1998; MacDonald et al., 2006) or the “foot” signal observed by amperometry (Braun et al., 2007), indicating that they reflect separate mechanisms or pore regulation. Indeed, we show here that PI(4,5)P_2_ transients delay peptide release (i.e. restrict late pore expansion), and that they coincide with the recruitment of an endocytic fission machinery at a time when the SNARE fusion machinery has already collapsed (Gandasi et al., 2015). Taken together, this supports the idea that the fusion reaction rapidly passes through a reversible state that is accelerated by PI(4,5)P_2_, after which pore expansion is relatively slow and can be further restricted by PIP(4,5)P_2_ dependent recruitment of an endocytic fission machinery. Although not studied here, this mechanism may in the extreme lead to endocytosis of largely intact granules by cavicapture (Obermuller et al., 2005; Taraska et al., 2003; Tsuboi et al., 2004).

Both PI(4)P and PI(4,5)P_2_, but not PI(3,4,5)P_3_, appeared transiently at the exocytosis site. Since interfering with PI4K or PIP5K blocked this local accumulation, it is likely the result of de novo synthesis by sequential action of these PI modifying enzymes. PI4K activity is found on granules and synaptic vesicles (Guo et al., 2003; Schvartz et al., 2012; Wiedemann et al., 1996). One possibility is therefore that PI(4)P, which is enriched in the granule membrane (non-zero ΔF/S in Fig 1E,H and (MacDonald et al., 2015), becomes available as substrate for the plasma membrane-localized PIP5K when the two membranes merge during exocytosis, which leads to conversion of PI(4)P to PI(4,5)P_2_ and a switch in membrane identity during exocytosis. Since blocking PI5K function prevents PI(4,5)P_2_ transients, it is unlikely the result of a diffusion-driven process in which the PI is sequestered by protein-lipid interaction (e.g., amphiphysin) (Picas et al., 2014). Notably, we did not detect any accumulation of PI(4,5)P_2_ before the moment of fusion (Omar-Hmeadi et al., 2018), which is in contrast with previous work (Aoyagi et al., 2005; Bogaart et al., 2011; James et al., 2008; Murray and Tamm, 2011). As discussed elsewhere, one reason for this may be the use of live cells rather than membrane sheets or model membranes (Omar-Hmeadi et al., 2018).

The neck of the vesicle likely contributes to accumulation of PI(4,5)P_2_ by hindering lipid diffusion during the early stages of fusion. This is suggested by the lack of FM4-64 efflux before full fusion (Fig 2C), and could maintain the vesicle membrane as isolated reaction environment in which PI5K activity leads to accumulation of PI(4,5)P_2_. This idea is consistent with superresolution measurements in which a PI(4,5)P_2_ increased in the vesicle membrane after fusion of the two membranes (Shin et al., 2018), although the notion of local synthesis was not tested in this work. Retention of PI(4,5)P_2_ may also occur as result of co-clustering of PI-binding proteins in microdomains, as has been shown *in vitro* for yeast BAR domain proteins (Zhao et al., 2013). In an alternative scenario, PI may cross the fusion pore, to be converted to PI(4,5)P_2_ by sequential action of PI4K and PI5K. However, this is less likely because interfering with PI4K activity reduced PI(4)P while having only minor effects on fusion pore behavior and PI(4,5)P_2_ generation, suggesting that PI(4)P serves mainly as a precursor for PI(4,5)P_2_. Regardless of the details, the mechanisms are likely different from the early stages of clathrin mediated endocytosis, where PI(4,5)P_2_ is generated in the plasma membrane by a positive feedback of enzymatic action and recruitment of PI5K, but does not accumulate in the vesicle membrane (Krauss et al., 2006).

PI(4,5)P_2_ binds to PH, PX, ENTH, and also to positively charged BAR domains, which likely mediate the recruitment of amphiphysin, amisyn, endophilin, epsin, and SNX9. Support for this comes from the strong correlation between the PI(4,5)P_2_ transients and protein recruitment (Fig 4), but it should be noted that amphiphysin, endophilin and SNX9 also contain curvature sensing domains that may be involved in their specific binding to the highly curved neck structure (Boucrot et al., 2012). Although several of these proteins are involved in CME, PI(4,5)P_2_ driven fusion pore regulation must occur by a different mechanism because clathrin was absent, and because genuine CME events were observed in parallel and exhibited a vastly slower timecourse. For most of the tested proteins the timecourse, onset, and amplitude of their recruitment mirrored that of the PI(4,5)P_2_ signal, with dynamin being a notable exception. Dynamin is key to the fission reaction of CME by forcing the GTP-hydrolysis dependent constriction of the neck of the budding vesicle, which promotes scission of endocytic vesicles. Dynamin is known to be involved in kiss&run exocytosis (Graham et al., 2002; Shin et al., 2018; Tsuboi et al., 2004), and there is support for amphiphysin-dependent recruitment of dynamin to the release site (Somasundaram&Taraska MBoC 2018). Dynamin mutants lacking GTPase activity affect peptide release kinetics in INS1 cells (Trexler et al., 2016), which suggests that fusion pore behavior ultimately depends on active dynamin. There is also evidence that proteins involved in membrane fusion, in particular the Ca^2+^-sensor synaptotagmin that binds PI(4,5)P_2_ via its C2B domain, can affect pore behavior by inducing local membrane curvature. However, synaptotagmin primarily affects transmitter release kinetics (Wang, 2001) and the mechanism is therefore distinct from the long-lasting pore restriction we observe here. It is also unlikely that fusion pore behavior is regulated by PI(4,5)P_2_-dependent actin polymerization around the vesicle, as for example during the slow fusion of much larger *Xenopus* egg cortical granules (Sokac et al., 2003), because actin is excluded from the insulin granule release site (Trexler et al., 2016)).

An importing remaining question is why PI(4,5)P_2_ accumulation and fusion pore restriction occurs only in a subset of granules. While it is possible that features on either granule or the release site predetermine the outcome of a particular fusion event, the continuous distribution of the pore lifetimes rather suggests a stochastic process where random variation in e.g. the local concentration of lipids, kinases and effector proteins leads to a graded outcome. Indeed, we noticed that after block of PI5K some of the remaining few exocytosis events with long fusion pore lifetimes had PI(4,5)P_2_ transients similar to control and recruited amphiphysin. Cellular signaling (e.g., cAMP/Epac), disease related protein over-expression, and perhaps location in polarized cells, will shift this continuum between full fusion and kiss&run exocytosis and thereby tune the secretory output to the cells demand for intercellular communication.

## Acknowledgments

We thank Per-Eric Lund for helpful comments, Olof Idevall for plasmids and helpful suggestions, and Jan Saras for plasmids and excellent technical assistance.

## Author contributions

M.O.H., A.G and S.B. designed experiments, analyzed the data, wrote, and reviewed the manuscript. M.O.H. performed experiments with PI sensors. A.G. performed experiments with proteins.

## STAR Methods

### Cells

INS1-cells clone 832/13(Hohmeier et al., 2000) were maintained at 37°C 5% CO_2_ in RPMI 1640 (Life Technologies) supplemented with 10% fetal bovine serum (FBS), streptomycin (100 U/ml), penicillin (100 U/ml), sodium pyruvate (1mM), HEPES (10mM), L-glutamine (2mM), and β-mercaptoethanol (0.5 mM). Cells were plated on coverslips coated with poly-L-lysine (Sigma) and transfected with plasmid DNA using Lipofectamine 2000 (Life Technologies) in Opti-mem (Life Technologies). Imaging was performed 24-36 hours after transfection.

Islets from diceased human donors were isolated and provided to us by the Nordic Network for Clinical Islet Transplantation (Uppsala, Sweden). All procedures were approved by the regional ethics committee EPN (Uppsala, Sweden). Islets were cultured free-floating in sterile dishes in CMRL 1066 culture medium containing 5.5 mM glucose, 10% fetal calf serum (FCS), 2 mM L-glutamine, streptomycin (100U/ml), and penicillin (100 U/ml) at 37°C in an atmosphere of 5% CO_2_ up to two weeks. The islets were then dispersed into single cells in 2 mL cell dissociation buffer (Thermo Fisher Scientific) supplemented with trypsin (0.005%, Life Technologies) followed by gentle pipetting for 30 seconds. The trypsin was then inhibited by adding 4 mL serum-containing medium and centrifuged for 5 minutes at 160 g. The resuspended cells were plated onto 22-mm poly-L-lysine coated coverslips and allowed to settle overnight. Cells were transfected with DNA plasmids using Lipofectamine 3000 (Life Technologies) in Opti-mem (Life Technologies) and infected using Adenovirus NPY-mCherry (Meur et al., 2010) or NPY-tdmOrange2 (Guček et al., 2019). Cells were imaged 24-36 hours post transfection and infection.

### Plasmids

PI(4,5)P_2_ was detected using the EGFP- and mRFP-tagged PH domains of PLCδ1 (PH-PLCδ1-EGFP or PH-PLCδ1-mRFP (Stauffer et al., 1998)). PI(4)P was detected using the EGFP-tagged P4M domain from *L. pneumophila* SidM (GFP-P4M-SidM (Hammond et al., 2014). PI(3,4,5)P_3_ was detected using the EGFP-tagged PH domain of the ARF protein exchange factor GRP1 (GFP-GRP1 (Idevall-Hagren et al., 2012)). Granule markers were NPY-EGFP, NPY-mCherry (Barg et al., 2010) and NPY-tdmOrange2 (Gandasi et al., 2015). CIBN-CAAX, GFP-CRY2-5ptaseOCRL, and CRY2-5ptaseOCRL were kindly provided by O Idevall(Idevall-Hagren et al., 2012), amphiphysin1-mCherry, GFP-CLC, endophilin2-mCherry, epsin2-mCherry and mCherry-SNX9 by CJ Merrifield (Taylor et al., 2011), dynamin1-GFP by W Almers (Merrifield et al., 2002), and mCherry-amisyn by the authors (Guček et al., 2019).

### Transient knockdown using siRNA or mirRNA

Transient Pip5k1c knockdown (KD) was performed using scrambled control siRNA or with predesigned siRNAs targeting the rat Pip5k1c sequence 5’-GCUACUAUGAACCUCAAtt-3’ (Ambion Life Technologies). Transfections were performed with Lipofectamine RNAiMAX (Life Technologies) according to the manufacturer’s instructions with a final siRNA concentration of 10 nM. For KD of PI4k2a a microRNA approach was used. A Mir30a-based cassette, targeting bases 499-519 (AAGAGCCATACGGGAACCTTA) of NM_053735.2 Rattus norvegicus phosphatidylinositol 4-kinase type 2 alpha (Pi4k2a) mRNA, was inserted between the stop-codon and the polyA signal in the expression plasmid NPY-mCherry(N1) (Barg et al., 2010). KD was routinely confirmed by quantitative RT-PCR.

### Quantitative RT-PCR

mRNA from control and Pip5k1c KD INS1 cells were extracted using NucleoSpin RNAPlus kit (Macherey-Nagel). RT-PCR were performed using QuantiTect SYBR Green RT-PCR kit (Qiagen) with the following primers: PIP5k1c-fwd2, 5’ GCAACCTCAGCTCTAAGCCA 3’; PIP5k1c-rev2, 5’ GGCAACAGGTGCATAGGTCT 3’; PIP5k1c-fwd1, 5’ AGACCTATGCACCTGTTGCC 3’; PIP5k1c-rev1 AATGAGTGGCTCGTTGCAGA 3’; PI4k2a-fwd1, 5’ GATCCTGGTTTCGACAGGGG 3’; PI4k2a-rev1, 5’ CAACAATCACGGGTGGCATC 3’; PI4k2a-fwd2, 5’ AGCCTGGTGGACCAAAAACT 3’; PI4k2a-rev2, 5’ AAGAGCTGGAATGACCCGAC 3’; CyclopA-fwd, 5’ GGTGACTTCACACGCCATAA 3’; CyclopA-rev, 5’ CTTCCCAAAGACCACATGC 3’; Ppia-fwd, 5’ TGTTCTTCGACATCACGGCT 3’; Ppia-rev, 5’ GTGTGAAGTCACCACCCTGG 3’. PCR reactions were performed using Light Cycler 2.0 (Roche), and results are expressed as ΔΔCt, normalized to the expression in control samples (scramble siRNA) in transient and stable KD cells.

### Solutions

Cells were imaged in an extracellular (EC) solution containing (in mM) 138 NaCl, 5.6 KCl, 1.2 MgCl2, 2.6 CaCl2, 3 D-glucose and 5 HEPES (ph7.4 adjusted with NaOH). For exocytosis experiments, glucose was increased to 10 mM and the solution was supplemented with 2 μM forskolin and 200 μM diazoxide, a K^+^-ATP-channel opener that prevents glucose-dependent depolarization. Exocytosis was then evoked with high K^+^ solution (75 mM KCl equimolarly replacing NaCl) pressure applied through a glass pipette similar to those used for patch-clamp experiments. All experiments were carried out at 37°C. 300 μM of BODIPY^®^ FL Phosphatidylinositol 4,5-bisphosphate (BODIPY FL-PI(4,5)P_2_; Echelon Biosciences, Salt Lake City, USA), was freshly prepared and incubated with 100 μM Histone H1 Carrier (Echelon Biosciences, Salt Lake City, USA) for 10 min at room temperature after vigorous pipetting to facilitate complex formation (Weiner et al., 2002). It was further diluted 1:10 with Hanks buffered saline solution. 150μl of the prepared complexes was dropped onto confluent INS1 cells in dark at 37 °C. After 20 minutes of integration, the medium was removed and replaced with a fresh EC solution. The addition of histone H1 alone served as a negative control. Kinase inhibitor GSK-A1 was used at concentration of 100nM (SYNkinase, Australia). To non-specifically stain the plasma membrane, styryl dye FM 4-64 (Thermo Fisher Scientific) (5 μg/mL) was puffed on the cell 30s before and during the K^+^-stimulated exocytosis using a double barrel pipette.

### Microscopy

Cells were imaged using a custom-built lens-type total internal reflection (TIRF) microscopy based on an Axio Observer Z1 with a 100x/1.45 objective (Carl Zeiss). Excitation was from two DPSS lasers at 491 and 561 nm (Cobolt, Stockholm, Sweden) passed through a cleanup filter (zet405/488/561/640x, Chroma) and controlled with an acousto-optical tunable filter (AA-Opto, France). Excitation and emission light were separated using a beam splitter (ZT405/ 488/561/640rpc, Chroma). The emission light was chromatically separated onto separate areas of an EMCCD camera (Roper QuantEM 512SC) using an image splitter (Optical Insights) with a cutoff at 565 nm (565dcxr, Chroma) and emission filters (ET525/50m and 600/50m, Chroma). Scaling was 160 nm per pixel. For movies, cells were excited simultaneously with 491 and 561 nm light and recorded in stream mode with 100 ms exposures (10 frames*s^−1^). The alignment of the two-color channels was corrected as previously described (Taraska et al., 2003).

Fusion pore experiments in INS1 cells and with A1 were imaged using a custom-built lens-type TIRF microscope based on an AxioObserver D1 microscope and a 100x/1.45 NA objective (Carl Zeiss). Excitation was from two DPSS lasers at 473 nm and 561 nm (Cobolt), controlled with an acousto-optical tunable filter (AOTF, AA-Opto) and using dichroic Di01-R488/561 (Semrock). The emission light was separated onto the two halves of a 16-bit EMCCD camera (Roper Cascade 512B, gain setting at 3,800 a.u. throughout) using an image splitter (DualView, Photometrics) with ET525/50m and 600/50m emission filters (Chroma). Scaling was 100 nm per pixel.

### Image analysis

Fluorescence intensities were measured as average fluorescence per pixel in the following regions of interest: (1) An area encircling a cell of interest (*cell*); (2) a background area outside the cell’s footprint (*bg*). Background-subtracted measurements are denoted by the variable *F=cell-bg*; (3) circles (*c*) of 3 pixel diameter centered within one pixel of the locations of solitary granules or, more rarely, of PI(4,5)P_2_ transients; (4) in an annulus (*a*) surrounding the circle with a diameter of 5 pixel. The background corrected annulus value was *S=a-bg* and reports local protein expression that is not related to the presence of a granule. We define Δ*F=c-a*, which estimates the fluorescence that is specifically localized to a granule (or in some cases a PI(4,5)P_2_ transient). We then calculate Δ*S=ΔF/S*, which normalizes for local expression level and is proportional to the apparent affinity of the measured protein to the granule site (Barg et al., 2010). For our object based analysis, we selected granules (usually in the red channel) that appeared diffraction-limited and round, were not located at the edge of the cell and separated from their neighbors. Analysis of the corresponding locations in the green channel was carried out automatically using Metamorph journal functions and Excel. Granules that were less than 1μm from the edge of the cell were excluded. So were the granules in Human islet preparations that were spatially and temporarily so aligned, they could not be separated. Exocytosis events were detected manually based on sudden disappearance of the granule fluorescence (first significant change ie. approx. 2 s.d. from the pre-exocytosis baseline). Protein accumulation at the PI(4,5)P_2_ transients site was aligned to the first rising PI(4,5)P_2_ frame and quantified. The PI(4,5)P_2_ (Fig 1–3) or protein binding (Fig 4) Δ*F/S* was calculated as a difference between the average of 5 frames at the peak of the single cluster, devided by the average of 5 frames at the beginning. Unless otherwise specified, images are displayed linearly auto-scaled based on their minimum and maximum brightness.

### Statistics

Experiments were repeated with at least 3 independent preparations. Data are presented as mean±s.e.m. unless otherwise stated. The violin plot outlines were generated in OriginLab 2019 (0% extended bandwidth) and illustrate kernel probability density, i.e. the width of the shaded area represents the proportion of the data located there. Boxplots indicate the median and quartiles with whiskers reaching up to 1.5 times the interquartile range. Statistical significance was assessed in OriginLab 2019 using Students t-test for two-tailed, paired or unpaired samples, one-way ANOVA, Mann-Whitney u-test, or Kolmogorov-Smirnov (KS) test, as appropriate and indicated in the figure legends. Significant difference is indicated by P-value next to the data.

