## Supplementary figures for "Local PI(4,5)P_2_ generation controls fusion pore expansion during exocytosis"

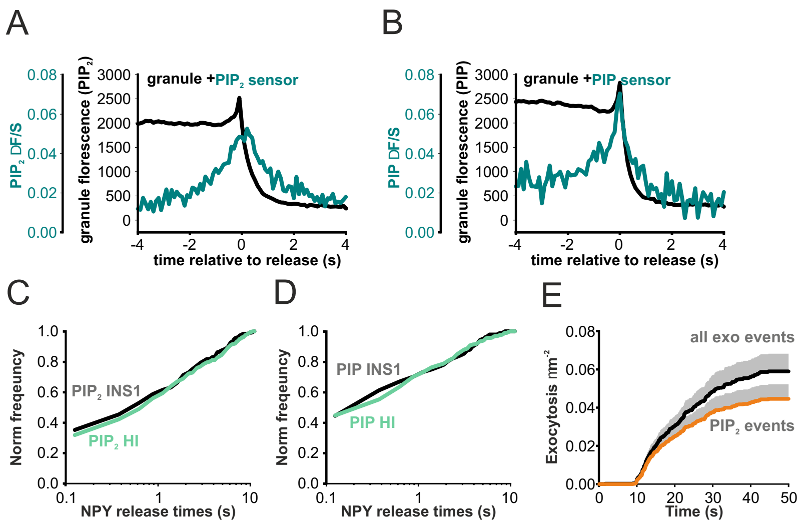


**Fig S1, related to Figure 1: Timecourse and effect on release time of EGFP-PH PLCδ1 and EGFP-P4M-SidM**

Additional analyis of data in Fig 1D-H.

(A) Average time course of NPY-tdmOrange2 (black) and the PI(4,5)$P_2$ sensor EGFP-PH-PLCδ1 fluorescence during exocytotic events (247 events in 26 INS1-cells). Same data as in Fig 1D, but aligned to moment of release.

(B) Average time course of NPY-tdmOrange2 (black) and the PI(4)P sensor EGFP-P4M-SidM fluorescence during exocytotic events (191 events in 26 INS1-cells).Same data as in Fig 1E, but aligned to moment of release.

(C) Cumulative frequency histograms of NPY release times in INS1 cells (black) and human beta-cells (green). All cells co-expressed NPY tdmOrange2 and EGFP-PH-PLCδ1. Same data as in Fig 1D-G.

(D) As in A, but for cells co-expressing EGFP-P4M-SidM . Same data as in Fig 1E-H

(E) Time course of all NPY release events (gray) and for the subset during which the EGFP-PH-PLCδ1signal

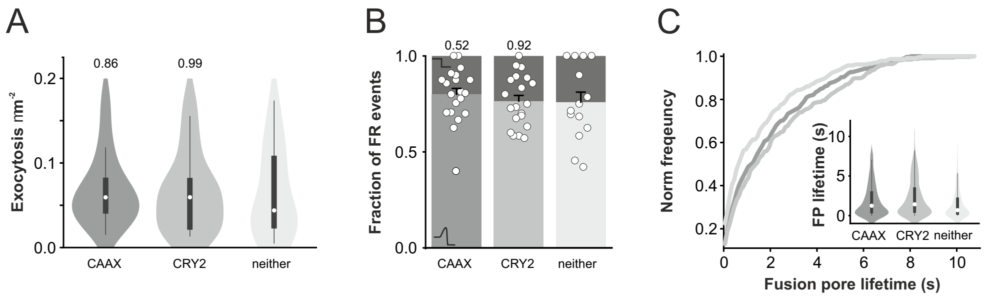


**Fig S5, related to Figure 5: Control experiments for the optogenetic experiment**

(A) Exocytosis during 40 s of $K^+$-stimulation for cells expressing NPY-EGFP with only CAAX (20 cells), only CRY2 (20 cells) or neither of them (14 cells), t-test

(B) Fraction of flash events as in A. Numbers indicate p-values, u-test

(C) Cumulative frequency histograms of NPY release times for same cells as in A. n=330 (CAAX), n=335 (CRY2) and n=236 (neither).

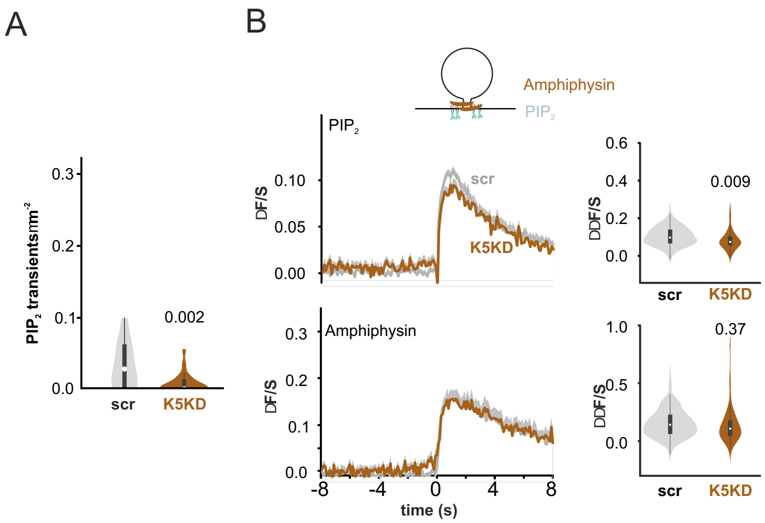


**Fig S6, related to Figure 6: Knock down of PIP5K1c decreases PI(4,5)$P_2$ transients**

(A) Normalized count of PH-PLCδ1-EGFP transients during 40 s of K+-stimulation in cells co-expressing PH-PLCδ1-EGFP and amphiphysin-mCherry, either with scrambled control (scr, 21 cells) or PIP5k1c specific shRNA (K5KD, 38 cells). Numbers on top of violins indicate p values, t-test

(B) Average time course (+SEM) of EGFP-PH-PLCδ1 (top) and amphiphysin-mCherry (bottom) in scrambled control (scr, 152 events in 21 cells) or K5KD, 62 in 38 cells). Violin plots depict distribution of peak amplitudes(DDF/S), non-paired t-test.

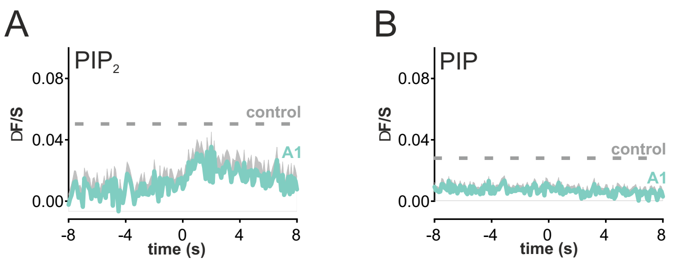


**Fig S7, related to Figure 7: Kinase-4 inhibitor (A1, 100nM) prevents PI(4)P, but not PI(4,5)$P_2$ transients in Human b cells.**

(A) Quantification of PI(4,5)$P_2$ (EGFP-PH-PLCδ1) accumulation at fusion site (ΔF/S, bottom) for the total number of events.

(B) Quantification of PI(4)P (EGFP-P4M-SidM) accumulation at fusion site (ΔF/S, bottom) for the total number of events.
